# Prediction and *in vitro* silencing of two lipid biosynthetic genes viz. *SCD* and *SREBP1* in chicken using RNA interference

**DOI:** 10.1101/2020.10.21.348482

**Authors:** N. Govardhana Sagar, A. Rajendra Prasad, R.N. Chatterjee, Pushpendra Kumar, Bharat Bhushan, P. Guru Vishnu, T. K. Bhattacharya

## Abstract

RNA interference is a widely used post transcriptional silencing mechanism for suppressing expression of the target gene. In the current study, five shRNA molecules each against *SCD* and *SREBP1* genes were designed after considering the parameters like secondary structures of shRNA constructs, mRNA target regions, GC content and thermodynamic properties (ΔG overall, ΔG duplex and ΔG break-target) to knockdown their expression. After successful transfection of these shRNA molecules into the chicken embryonic hepatocyte culture, expression of the target genes were monitored by real time PCR. Significant reduction (P<0.05) in the expression of *SCD* and *SREBP1* genes with minimal activation of immune response genes in hepatocytes was observed after transfecting the shRNA molecules into them. The shRNA constructs against *SCD* gene showed the knock down efficiency ranged from 20.4% to 74.2%. In case of shRNA constructs against *SREBP1* gene, they showed knock down efficiency ranging from 26.8% to 95.85%. These results clearly demonstrated the successful down regulation of the gene expression by designed shRNA molecules against both the target genes under *in vitro* condition. The shRNA2 molecule for *SCD* gene and the shRNA1 molecule for *SREBP1* gene were found as the best among all the shRNA molecules used for silencing the target genes under cell culture system. It is concluded that the shRNA molecules designed against SCD and SREBP1 genes showed potential to silence the expression of these genes under in vitro chicken embryonic hepatocyte cell culture system.

## Introduction

Stearoyl-CoA desaturases (*SCD*), an endoplasmic reticulum-resident enzyme catalyzes introduction of the first double bond in the *cis*-delta-9 position of several saturated fatty acyl-CoAs, principally palmitoyl-CoA and stearoyl-CoA, to yield palmitoleoyl- and oleoyl-CoA, respectively [1, 2]. Stearoyl-CoA desaturase (*SCD*) is an iron-containing enzyme that catalyzes a rate-limiting step in the synthesis of unsaturated fatty acids. The principal product of *SCD* is oleic acid, which is formed by desaturation of stearic acid. The ratio of stearic acid to oleic acid has been implicated in the regulation of cell growth and differentiation through effects on cell membrane fluidity and signal transduction.

Sterol regulatory element binding protein-1 (*SREBP1*) is a transcription factor that binds to a sequence in the promoter of different genes, called sterol regulatory element-1 (SRE1). This element is a decamer flanking the LDL receptor gene and other genes involved in sterol biosynthesis. *SREBP1* regulates genes required for glucose metabolism and fatty acid and lipid production and its expression is regulated by insulin [3]. *SREBP1* regulates genes related to lipid and cholesterol production and its activity is regulated by sterol levels in the cell [4; 5]. The transcription factor *SREBP1* regulates *de novo* lipogenesis in the liver in response to increases in insulin. *SREBP*s are transcription factors of the basic helix-loop-helix leucine zipper family that are synthesized as precursors and bound to the endoplasmic reticulum membrane. In the present study, we have developed an *in vitro* model for reduction of *SCD* and *SREBP1* in chicken embryonic liver cell culture by using the artificially synthesised shRNA molecules.

## Materials and Methods

### Experimental animals

The study was conducted in the control broiler chicken line maintained at ICAR-Directorate of Poultry Research, Rajendranagar, Hyderabad, India. The fertile eggs were collected from control broiler chicken and incubated at 98°F and 85% relative humidity in the incubator for 12 to 13 days. The embryonated eggs of 12-13 days old were taken out for preparation of embryonic hepatocyte primary cell culture. The whole experiment was approved by the Institute Animal Ethics Committee (IAEC) and Institute Bio-safety Committee (IBSC) of ICAR-Directorate of Poultry Research, Rajendranagar, Hyderabad, India to carry out the animal experiment.

### *In silico* designing of shRNA molecules

In the present study, pENTR /U6 Entry Vector (Invtitrogen) was used to facilitate the generation of an entry construct that permits high-level expression of shRNA of interest in eukaryotic cells for RNAi analysis of a target gene (Fig.S1). The pENTR/U6 vector contains a U6 cassette containing elements which is needed for allowing RNA Polymerase III - controlled expression of the shRNA of interest in eukaryotic cells and a cloning site containing 4-nucleotide 5’ overhangs on each DNA strand for directional cloning of the ds-oligo encoding the shRNA of interest. For designing of shRNA molecules, firstly DNA single stranded oligos that are targeting the Open Reading Frame (ORF) of *SCD* and *SREBP1* genes were designed by using the BLOCK-iT RNAi Designer (https://rnaidesigner.thermofisher.com/rnaiexpress/) software (Table 1).

**Table 1.**
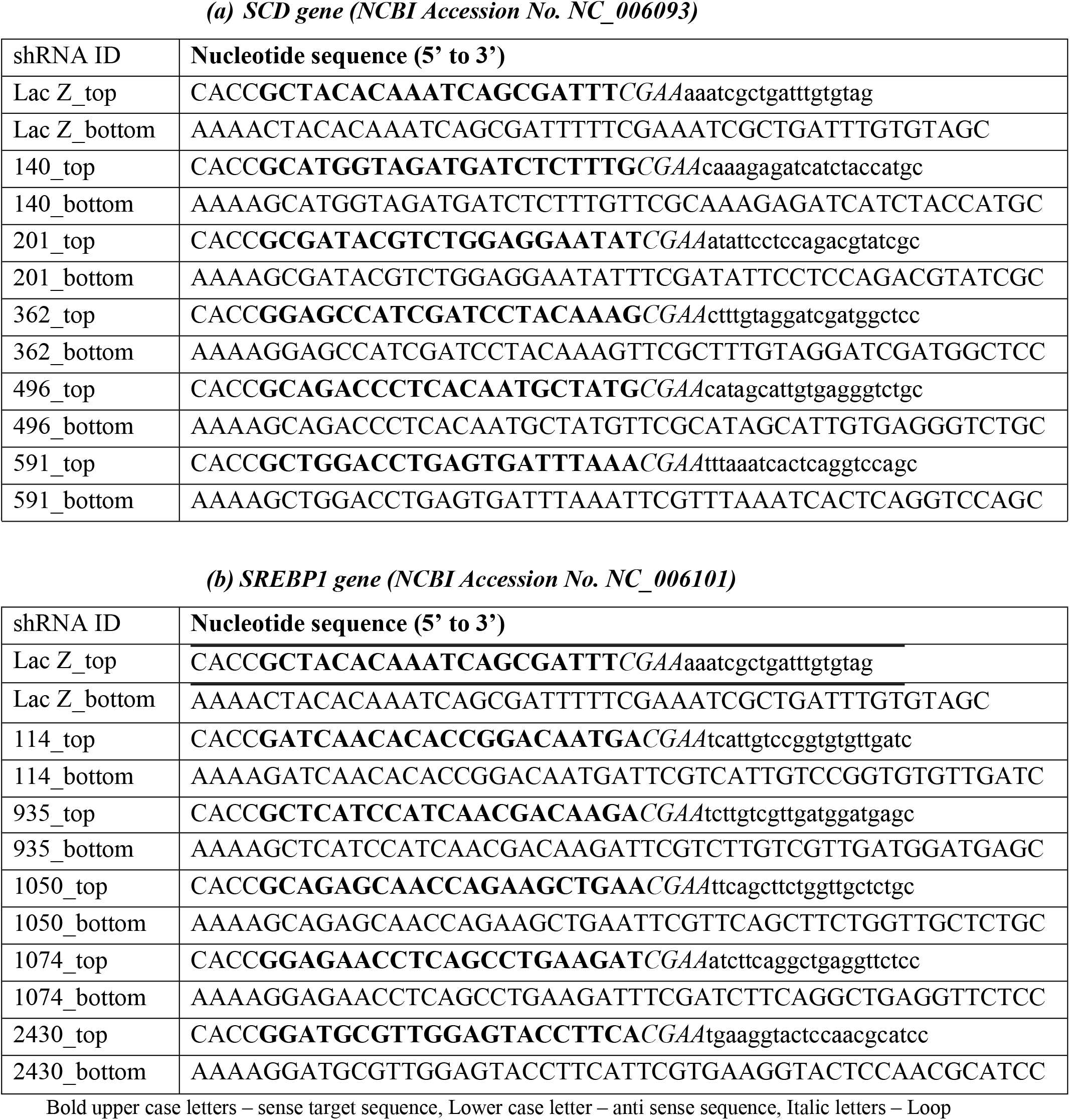
List of designed shRNA molecules against *SCD* and *SREBP1* genes.

A final concentration of 50 μM double stranded oligos were prepared by mixing top strand DNA oligo (200 μM) – 5 μl, Bottom strand DNA oligo (200 μM) - 5μl, 10X Oligo annealing buffer - 2μl and DNAse/RNase free water - 8μl, by using the conditions 95°C for 4 minutes and then allowed to cool at room temperature for 5-10 minutes to allow the single-stranded oligos to anneal. The sample was placed in a micro centrifuge and centrifuged briefly (~5 seconds) and mix gently.

### Prediction of secondary structures of *SCD* and *SREBP1* mRNA and shRNA molecules

For predicting the secondary structures for *SCD* and *SREBP1* mRNAs, web server of Mfold 2.3 version [6] (http://mfold.rna.albany.edu/?q=mfold/RNA-Folding-Form) was used after considering the all the required criteria. Secondary structures of antisense strands of shRNA molecules against *SCD* and *SREBP1* genes were predicted by using the RNA fold program of the Vienna RNA web service version 2.0 [7] (http://rna.tbi.univie.ac.at/cgi-bin/RNAWebSuite/RNAfold.cgi).

### Prediction of thermodynamic parameters of shRNA molecule and its target region

Oligowalk program [8] in the RNA structure version 5.3 (http://rna.urmc.rochester.edu/cgi-bin/server_exe/oligowalk/oligowalk_form.cgi) was used to predict the thermodynamic properties that governs the binding affinity of shRNA molecules and its target region on mRNA molecules [9]. Following thermodynamic properties [10] were included in the present study:

i. **<u>ΔG overall</u>**: The net energy (ΔG in kcal/mol) resulting due to binding of target site by oligos, after consideration of all energy contributions viz. oligo-self-structure energy and target structure breaking energy. The higher negative value of ΔG signifies stronger binding.
ii. **<u>ΔG duplex</u>**: It determines the stability of the duplex formed between guide strand of siRNA and the nucleotides at the target site. A less negative value of ΔG duplex denotes less stability of duplex and vice versa.
iii. **<u>ΔG break−target (disruption energy)</u>**: The energy cost for disrupting local secondary structure at the mRNA target region. A more negative value denotes that the binding site is not completely open and less accessible.

### Cloning of ds oligos into pENTR /U6 entry vector

The ds oligos (50 μM stock) were diluted to a final concentration of 5 nM by performing two 100-fold serial dilutions. First 100-fold dilution was accomplished with DNase/RNase-free water and then second100-fold dilution was completed with 1X Oligo annealing buffer supplied with the kit. These diluted ds oligos were used for cloning into the pENTR /U6 Entry Vector. The annealed shRNA oligo nucleotides were ligated into RNAi Ready pENTR™ U6 vector by preparing 20 μl of ligation mixture containing 5X Ligation Buffer - 4 μl, pENTRTM /U6 (0.5ng/ìL) - 2 μl, ds oligo (5 nM; 1:10000 dilution) - 1 μl, T4 DNA Ligase (1 U/μL) - 1 μl and DNAse/RNase free water - 12 μl. This ligation mixture was mixed well and incubated for 30 minutes at room temperature. Scrambled shRNA oligos (lacZ) supplied with the kit were also ligated to pENTR™/U6 vector and used as a negative control in the experiment.

Recombinant vector containing the shRNA molecules were transformed into the One Shot^®^ TOP10 chemically competent *E.coli* cells by giving heat-shock (42°C for 30 seconds and then, immediately transferred to ice). Transformed cells were grown in 250μl of super optimal broth with catabolite repression (S.O.C.) medium at room temperature by incubating in horizontal shaker incubator (200 RPM) at 37°C for 1 hour. 200 μL of transformation mixture was spread on a pre-warmed LB agar plate containing 50 μg/ml of kanamycin and incubated overnight at 37°C.

Kanamycin resistant colonies were picked and inoculated into the LB broth containing kanamycin 50 μg/mL of media and incubated over night at 37°C to analyze the positive clones. Kanamycin resistant clones were screened for the presence of shRNA in the plasmid constructs by performing colony PCR with U6 forward primer: 5’GGACTATCATA TGCTTACCG3’ and M13 reverse primer: 5’-CAGGAAACAGCTATGAC-3’ by using 2.5 μL of 10XPCR buffer, 0.5 μL of dNTP mix (2.5 mM), 1.5 μL (30 ng) each of forward and reverse primers, 0.2 μL (1U) of Taq DNA polymerase, 2 μL of colony lysate and nuclease free water to make the volume up to 25 μL in 0.2 ml PCR tubes. Thermal cycling conditions followed were, initial denaturation at 95°C for 10 minutes followed by 35 cycles of denaturation at 95°C for 30seconds, primer annealing of 54°C for 30 seconds and extension at 72°C for 30 seconds and final extension of 72°C for 10 minutes.

Plasmids containing shRNA molecules against *SCD* and *SREBP1* genes were isolated by using Gene JET Plasmid Miniprep Kit (#K0503, Thermo Scientific, USA). The plasmid obtained from each pENTR™/U6 entry construct was sequenced to confirm the sequence and correct orientation of the ds oligo insert.

### Establishment of primary chicken embryonic hepatocyte (CEH) culture

Primary chicken embryonic hepatocyte culture was established by using 12-13 days old embryos. Embryos were collected aseptically by breaking the broad end of the egg after piercing the chorio-allantoic membrane (CAM) and placed in 9 cm petri-dish containing sterile Phosphate Buffer Saline (PBS) and rinsed thoroughly. Head, limbs and wings were separated and then, ventral side of the embryo was cut opened for collecting the liver lobes into the petri-dish containing the PBS. After thorough washing in PBS and mincing, liver tissues were transferred into the beaker containing sterile magnetic bar and approximately 10-15 ml of 0.125% trypsin. The beaker was placed on the magnetic stirrer for stirring at 37°C at about 100 RPM for less than 10 minutes. After allowing the pellets to settle down, supernatant was filtered through sterile double layered muslin cloth into a fresh beaker. The filtrate was centrifuged for 5 minutes at 3000 RPM and to stop the trypsin action, the resulting pellet was re-suspended in 5ml of growth medium (DMEM, Sigma) containing Fetal Bovine Serum (FBS). Now, the media with the liver cells were centrifuged at 3000 RPM for 3 min and then, the resulting pellet was re-suspended in 5ml of growth medium containing 10% FBS, 1% Tryptone Phosphate Broth and antibiotic-antimycotic solution. Hemocytometer was used to count the number of cells and accordingly, the cell suspension was diluted to get a cell concentration of 1 × 10^6^ cells/ml. Approximately 2 × 10^5^ cells/ cm^2^ were seeded into the 25 cm^2^ tissue culture flask and incubated at 37°C with 5% CO_2_. Medium was changed at regular intervals to have good growth of cells and counter the depression of pH.

### Transfection of shRNA constructs into CEH Culture cells

All the shRNA recombinant plasmid constructs against *SCD* and *SREBP1* genes were transfected into the chicken primary embryonic hepatocyte culture in order to assess the activity of shRNA molecules in hepatocytes. Approximately 0.4 ml of the hepatocyte cell suspension and plasmid containing shRNA molecules against *SCD* and *SREBP1* genes were taken into electroporation cuvette and mixed gently. Single square wave pulse was given at a voltage of 150 mV for pulse length of 10 milli second. Both, pENTR U6/lacZ shRNA and pcDNA 1.2/V5/lacZ reported plasmids were co-transfected in to CEH. The pcDNA 1.2/V5/lacZ reported plasmid is used as a positive control for RNAi response in CEH. For getting optimal results, we had used 6 fold more entry construct DNA than reporter plasmid during co-transfection. Immediately after transfection, approximately 200 μl of cell suspension transferred to each well of a well plate containing 1.8 ml of growth medium and incubated at 37°C with 5% CO_2_ level.

### RNA extraction and cDNA synthesis

The cells were harvested after 48 hours of transfection and RNA was isolated for transient RNAi analysis. The total RNA was isolated from the hepatic cells grown after successful transfection of the plasmids containing shRNA molecules against both target genes as per the manufacturer’s protocol using Trizol (Sigma). The RNA samples were treated with DNase I (Fermentas) for removal of possible genomic DNA contamination. cDNA was synthesized by using High-Capacity cDNA Reverse transcription Kit (Applied Biosystems, #4368814) in a final volume of 20 μL containing 10X Reverse Transcription (RT) buffer (2 μL), 10X RT random primers (2 μL), 100 mM of 25X dNTP mix (0.8 μL), RNase inhibitor (1 μL), MultiScribeTM Reverse Transcriptase (1 μL), Nuclease-free H2O (3.2 μL) and 1 ng RNA (10 μL). Reverse transcription was carried out in thermocycler (Mastercycler, Eppendorf, Germany) following the manufacturer’s instructions, which includes the incubation of reaction at 25°C for 10 minutes followed by 37°C for 2 hours and 85°C for 5 minutes. The resulted cDNA was stored at −20°C till further use.

### Real time quantitative PCR

The mRNA expression levels of *SCD*, *SREBP1* genes and immune responsive genes viz. Interferon-A (IFN-A) and Interferon-B (IFN-B) genes in the transfected hepatic cells were quantified by using thermal cycler Applied Biosystems® Step One Real-Time PCR (Life Technologies) with SYBR® Green JumpStart™ TaqReadyMix™. Glyceraldehyde 3-phosphate dehydrogenase (*GAPDH*) was used as an internal control for normalizing different amounts of input RNA (Table.2). The realtime PCR was performed for each sample in triplicates.

**Table 2.**
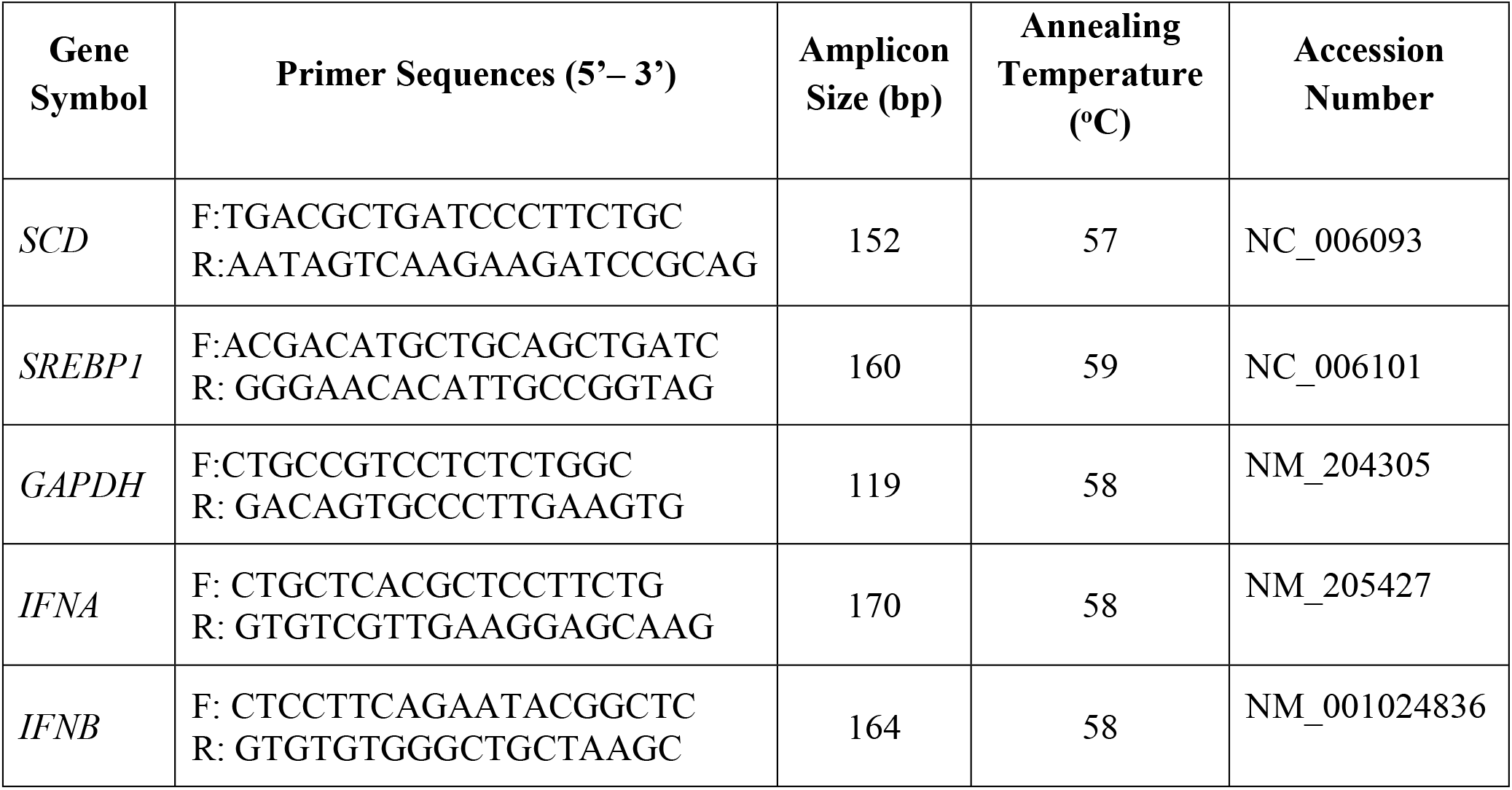
List of primers used for real time expression quantification of SCD, SREBP1 and immune response genes.

### Relative quantification

The gene quantification was expressed as “n-fold up/down regulation of transcription” in relation to a reference sample, called the calibrator (mock transfected hepatocytes). The expression of target gene was calibrated by that of the reference gene, *GAPDH*, at each time point and converted to the relative expression (fold of expression), as follows:

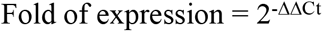

Where,

ΔC_t_ = Average C_t_ of target gene - Average C_t_ of reference gene (*GAPDH*)

ΔΔC_t_ = Average ΔCt of target (shRNA) – Average Δ C_t_ of calibrator sample (Mock)

The Knock down efficiency (KD%) of all the shRNA was calculated by considering scrambled shRNA as control.

### Statistical analysis

The experiments were repeated twice and statistical analysis was carried out using trial version of SPSS 20. Univariate General linear model with Tukey’s HSD and DMRT as post-hoc test was used to study the significant difference between different shRNA groups due to the knock down effect of target genes. Data from representative experiments were presented as Mean ± SE for different samples with differences determined by least significant differences at 5% level (P < 0.05). The degree of association between the expression of different genes was calculated by Pearson correlation coefficient.

### Production of hyper-immune sera for Western blotting and ELISA

For the production of hyper-immune sera against *SCD* and *SREBP1*, a 591 bp length of *SCD* cDNA and a 507 bp length of *SREBP1* cDNA were amplified by designing specific primers (SCD_Ab Forward: AAGCTTATGCACCACCATCACCATCATAATATCCTCATGAGCCTG; SCD_Ab Reverse: GGATCCAAACATGTGAGCGCTG; SREBP1_Ab Forward: AAGCTTATGCACCACCATCACCATCATCCTGACAGCACCGTGTC; SREBP1_Ab Reverse: GGATCCGTCTGCCTTGATGAAGTG). The forward primers of bothe cDNAs contained with 6X histidine tag and *BamHI* restriction enzyme while the reverse primers contained *Hind III* restriction enzyme. The PCR amplification was carried out in 200 μL PCR tubes containing 2.5 μL of 10X buffer, 1 μL of dNTP mix (2.5mM), 1.5 μL (30ng) each of forward and reverse primers, 0.3 μL (1.5U) of Taq DNA polymerase, 1 μL of genomic DNA and nuclease free water to make up the volume up to 25 μL. Thermal cycling conditions followed for both the genes were initial denaturation at 95 °C for 10 minutes followed by 35 cycles of denaturation at 95 °C for 30 seconds, primer annealing at 55 °C for 30 seconds and extension at 72 °C for 1 minute 30 sec and a final extension of 72 °C for 10 minutes. The amplified product was gel eluted and purified by using QIA quick gel extraction kit (Qiagen).

The pAcGFP1-C1 expression vector and purified amplicons of cDNA of both the genes were digested with *BamHI and Hind III* restriction enzymes. After RE digestion, cDNA of *SCD* gene and *SREBP1* gene were cloned separately into the pAcGFP1-C1 expression vector. The ligated products were transformed into DH5α *E.coli* competent cells. The positive clones were screened and identified by colony PCR, plasmid PCR and sequencing. The recombinant plasmids were isolated by using Gene JET plasmid miniprep kit (Thermo Scientific, USA). The recombinant plasmids of *SCD* and *SREBP1* genes were transfected into the CEH by using the Gene Pulser Xcell^TM^ Electroporation system (Biorad). After transfection, hepatocytes were grown in growth medium for 48 hrs and then, harvested for isolation of proteins. From the cell lysates, both *SCD* and *SREBP1* proteins tagged with 6x histidine were extracted by using the His-Spin Protein Miniprep^TM^ kit (GCC Biotech, Kolkata, India).

The purified protein was mixed with Freund’s Adjuvant (IFA) and the mixture was injected subcutaneously (s/c). Primary injection was given wit Freund’s Complete Adjuvant (CFA) and then, booster injections were given with Incomplete Freund’s Adjuvant (IFA). A total of 6 male Wistar rats (2 for *SCD* protein, 2 for *SREBP1* protein and 2 as control) of 8 weeks age were included in immunization schedule throughout period. The detailed Immunization protocol for hyper-immune sera production was as shown in Table 3. The IgG was purified from hyper immune sera using IgG purification kit (Himedia).

**Table 3.** Immunization protocol for rat polyclonal antibody production.

| **Procedure** | **Protocol Day** | **Description** |
| --- | --- | --- |
| Control serum collection | Day 0 | Pre-immune bleed (0.5ml per rat) |
| Primary Injection | Day 1 | Immunize with 0.1mg antigen in CFA, sc 4 sites |
| Booster Injections | Day 14 | Boost with 50µg antigen IFA, sc 4 sites |
| 2<sup>nd</sup> Booster Injections | Day 21 | Boost with 50µg antigen IFA, sc 4 sites |
| Test bleeds | Day 35 | Test-Bleed (0.5ml per rat) |
| 3<sup>rd</sup> Booster Injections | Day 42 | Boost with 50µg antigen IFA, sc 4 sites |
| Blood collection | Day 49 | Primary antibody production |

### SDS-PAGE and Western Blotting

A period of 48 h after transfection of chicken embryonic hepatocyte cells with shRNAs against *SCD* and *SREBP1* genes, cell pallets were harvested. The cell pellets were washed in ice cold PBS solution and then cell lysate was prepared by adding cold cell lysis buffer (150 mMNaCl, 1% NP-40, and 50 mM Tris, pH 7.4) @ 1ml per 150 cm^2^ flask, agitated constantly and centrifuged at a rate of 12,000 rpm, at 4 °C for 20 min. Equal volume of Laemmli 2X sample loading buffer (10% SDS, 0.025% Bromophenol blue and 1% DTT) added to the obtained cell lysate and boiled at 100 °C for 3 min. About 20 μl of digested samples containing approximately 15–20 μg of protein loaded into the wells of SDS-PAGE with discontinuous buffer system Tris–Glycine–SDS buffer, pH 8.3 to separate the protein mixture. After completion of electrophoresis, the gel containing protein was transferred on to the polyvinylidene fluoride (PVDF) membrane in the presence of Tris–Glycine–Methanol Buffer. After careful transfer of the gel, the blotted PVDF was immersed in 3% BSA blocking buffer with primary antibody (1:1000 dilution in TBST) and incubated at 4°C for overnight. Then the membrane was washed three times with the TBS Tween 20 (TBST) for 5 minutes for each wash. After washing, membrane was incubated with anti-rat IgG HRP conjugate (Sigma) diluted to 1:1000 in TBS Tween 20 buffer for 1.5 hours with constant agitation. After washings with the TBS Tween 20 (TBST), the PVDF membrane was incubated in DAB substrate solution for 5-30 minutes until the color development. Soon after the appearance of brown color, substrate solution was drained and the reaction was stopped by adding distilled water.

### Sandwich ELISA

IgGs specific for *SCD* and *SREBP1* have been pre-coated onto a 96-well plate (12 × 8 Well Strips) and blocked separately. Test samples (cell lysate obtained after transfecting the shRNA molecules into the hepatocytes) were added to the wells, incubated and removed. HRP detector antibody specific for *SCD* and *SREBP1* was added, incubated and followed by washing. HRP-Peroxidase Conjugate was then added, incubated and unbound conjugate was washed away. An enzymatic reaction was produced through the addition of TMB substrate which is catalyzed by HRP generating a blue color product that changes yellow after adding acidic stop solution. The density of yellow coloration read by absorbance at 450 nm is quantitatively proportional to the amount of sample *SCD* and *SREBP1* captured in well.

## Results

### Secondary structures of target mRNA regions and shRNA molecules

Local secondary structures of target mRNA sequences of *SCD* and *SREBP1* genes were predicted and they revealed that all the all the shRNA target mRNAs consists of stem and loop secondary structures (Fig.1 and Fig.2). Analysis of secondary structures of shRNA molecules designed against *SCD* and *SREBBP*1 genes revealed different types of secondary structures (Fig.3 and Fig.4).

**Fig 1.**
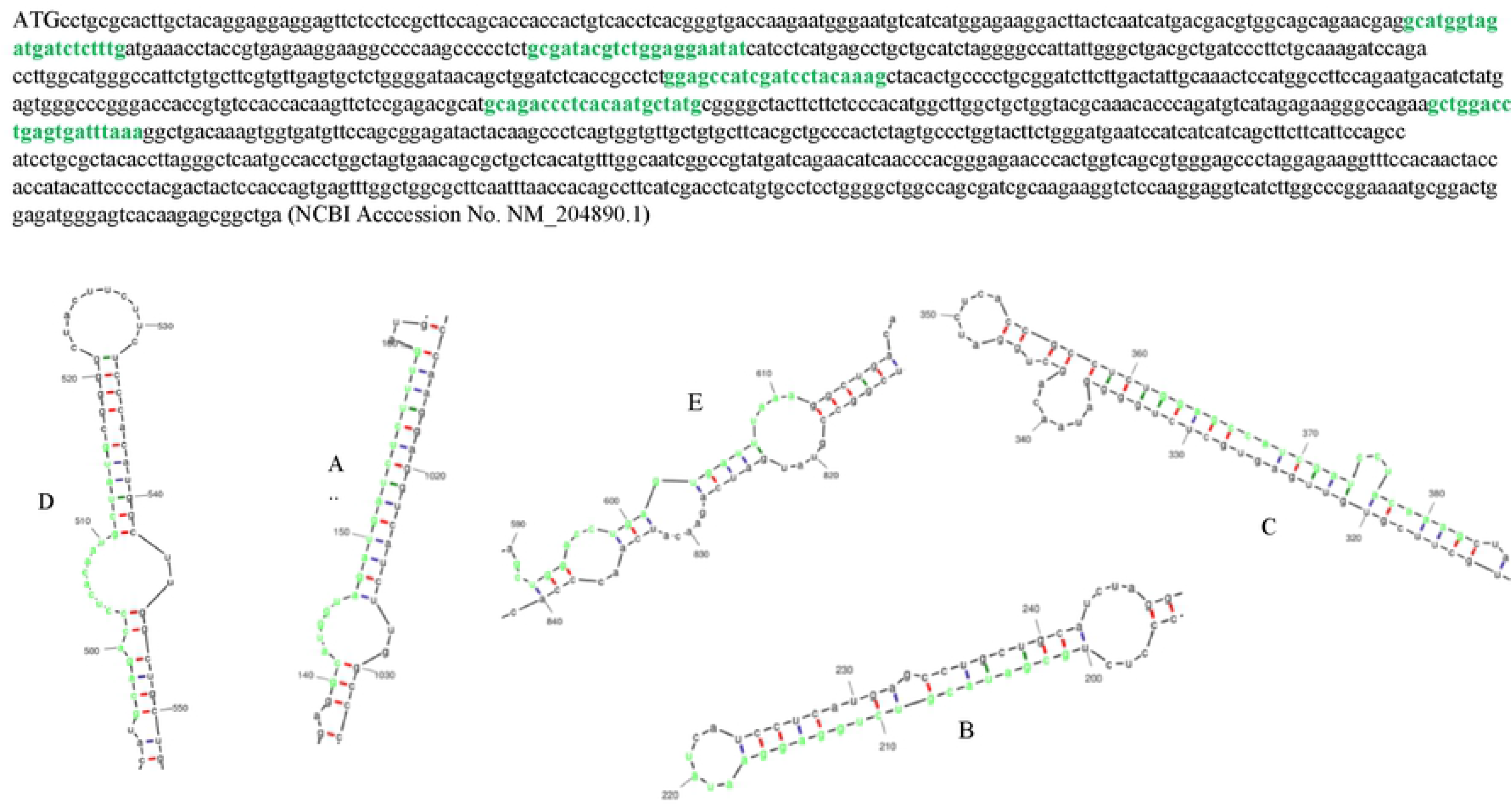
Local secondary structures of *SCD* mRNA targeted by shRNA I - A, shRNA2 - B, shRNA3 - C, shRNA4 - D and shRNA5 - E. Green highlighted region - mRNA region targeted by shRNA Red bar- most likely be single stranded, other colour bars: less likely to be single stranded.

**Fig 2.**
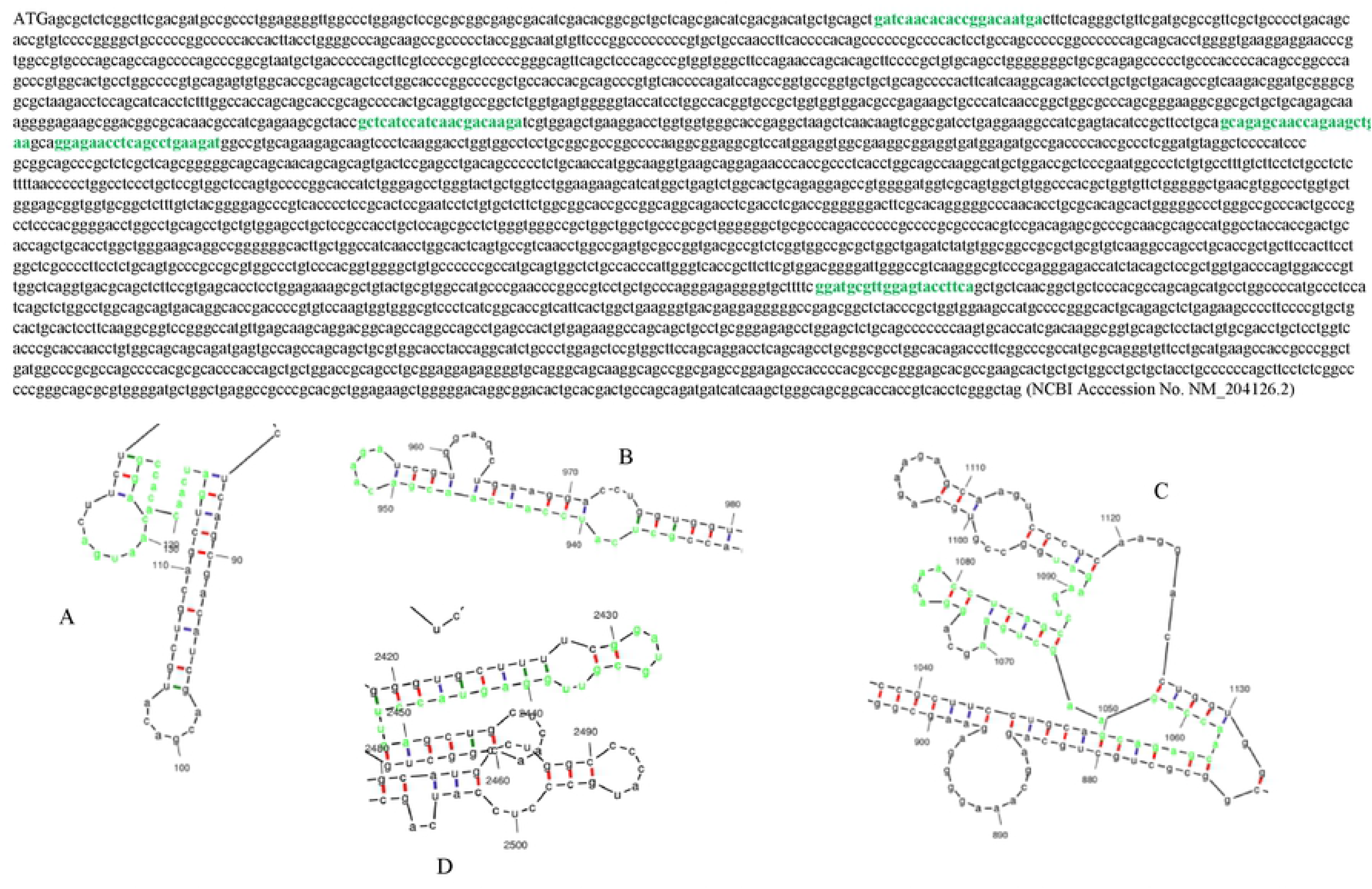
Local secondary structures of *SREBP1* mRNA targeted by shRNA1 - A, shRNA2 - B, shRNA3 & shRNA4 - C and shRNA5 - D. Green highlighted region - mRNA region targeted by shRNA. Red bar- most likely be single stranded, other colour bars: less likely to be single stranded.

**Fig 3.**
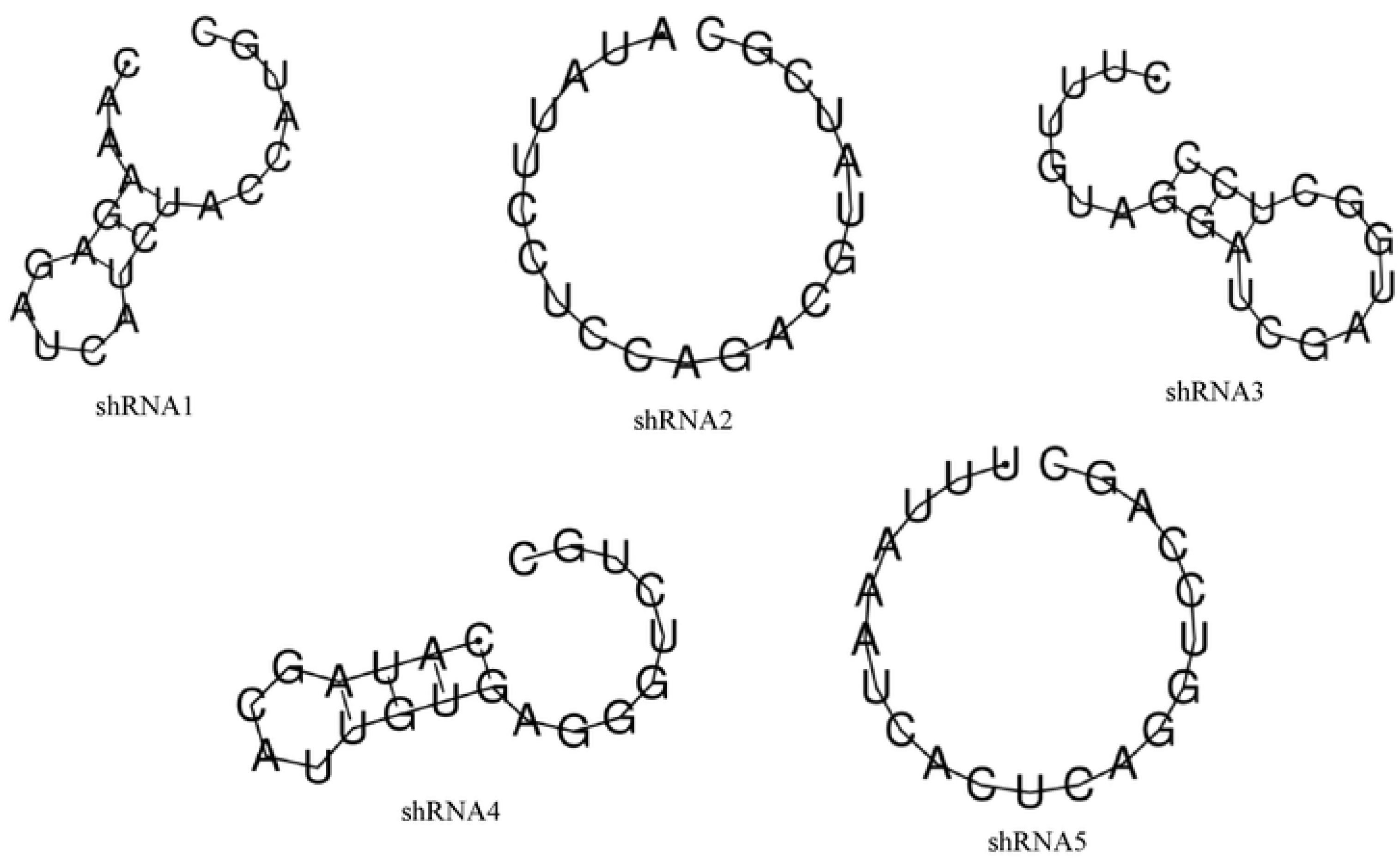
Secondary structures of anti sense/guide strand of different shRNA molecules designed against *SCD* mRNA.

**Fig 4.**
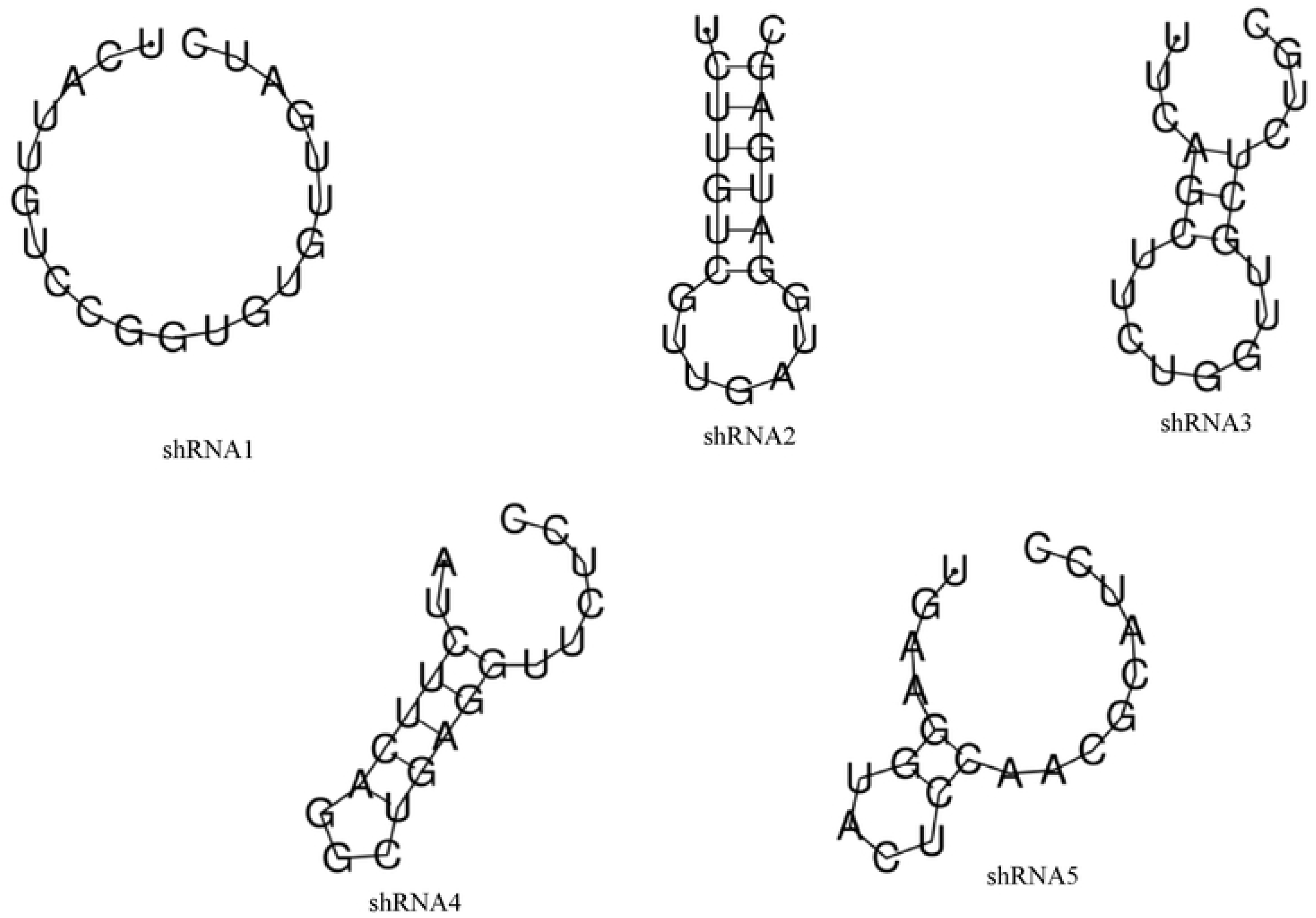
Secondary structures of anti sense/guide strand of different shRNA molecules designed against *SREBP1* mRNA.

### Thermodynamic properties

The predicted values (negative) of ΔG overall, ΔG duplex and ΔG break−target (disruption energy) for mRNA target regions of *SCD* and *SREBP1* genes were within the desirable range (Table 4 & Table 5). In case of *SCD* gene target mRNA region, overall ΔG value was almost same for shRNA2, shRNA4 and shRNA5. ΔG duplex value was the highest for shRNA5 and lowest for shRNA1. ΔG break−target (disruption energy) was highest for shRNA3, lowest for shRNA2. For *SREBP1* gene mRNA target regions, overall ΔG value was highest for shRNA1, lowest for shRNA2. ΔG duplex value was highest for shRNA4 and lowest for shRNA1. ΔG break−target (disruption energy) was highest for shRNA2 and lowest for shRNA1. GC content of the shRNA target mRNA regions of *SCD* gene ranged from 43% to 52%, whereas, GC content of the shRNA target mRNA regions of SREBP1 gene ranged from 48% to 52% (Table 4 & Table 5).

**Table 4.**
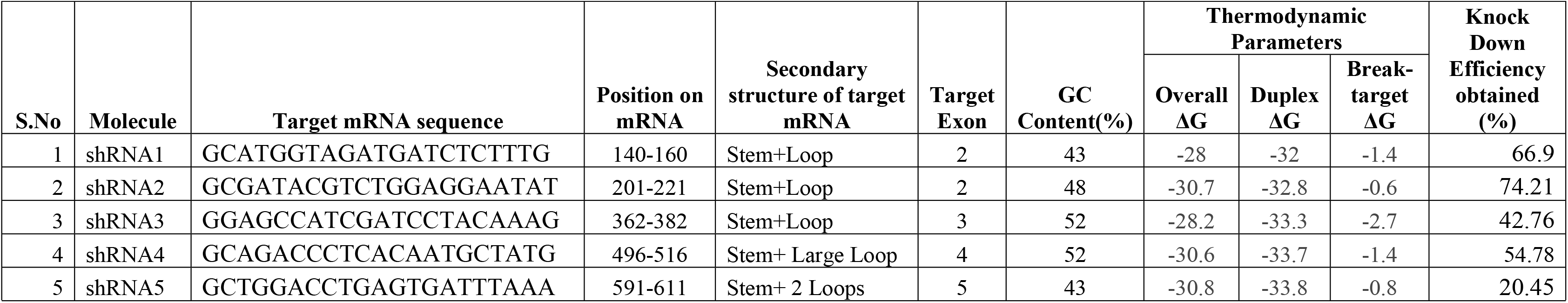
Thermo Dynamic properties & Knock Down Efficiencies of anti-*SCD* shRNA molecules.

**Table 5.**
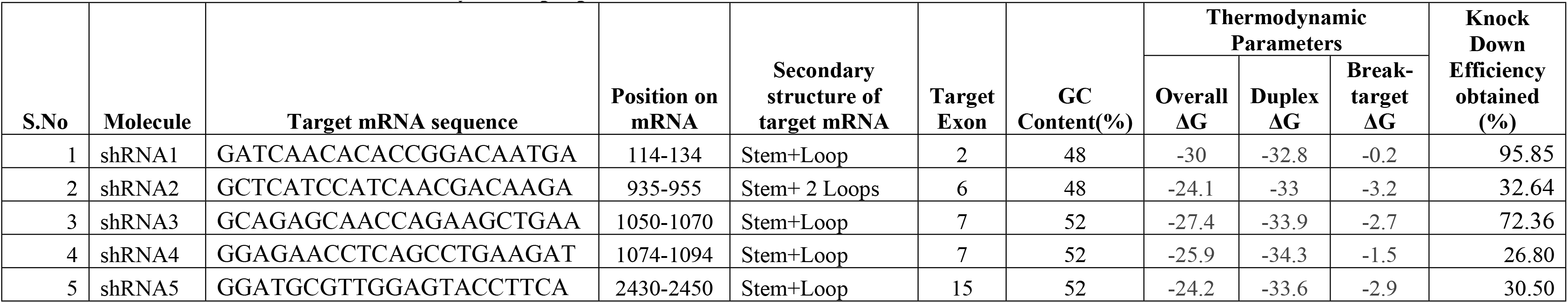
Thermo Dynamic properties & Knock Down Efficiencies of anti-*SREBP1* shRNA molecules.

A volume of 5 μl of 500 nM stock of ds oligos from each of the shRNA molecule was loaded on to the 4% agarose gel for checking the integrity of the annealed ds oligos (Fig S2 & Fig S3). Annealed ds oligos were observed at 50 bp length, unannealed single-stranded oligos were observed at 25 bp length as the agarose gel is non-denaturing. Therefore, the single-stranded oligos do not resolve at the expected size due to formation of secondary structure. Successfully annealed ds oligos were used further for cloning into the pENTR /U6 Entry Vector.

### Cloning of *SCD* and *SREBP1* anti-shRNA in pENTR/U6 Entry Vector

Successfully annealed ds oligos were used for cloning into RNAi-Ready pENTR/U6 entry vector. After successful ligation of annealed ds oligos into RNAi-Ready pENTR/U6 entry vector, recombinant clones were transformed into competent *E.coli* cells. Recombinant clones were confirmed by colony PCR (Fig S4 & Fig S5) and plasmid PCR (Fig S6 & Fig S7) on 1.5 % agarose gel and sequencing.

### Transient transfection of anti-*SCD* and -*SREBP1* shRNA construct into CEH

The hepatocytes were transfected with anti-*SCD* and -*SREBP1* shRNA constructs separately and grown in DMEM medium. The cells were harvested and then, total RNA and DNA were isolated from the harvested hepatocytes. For confirmation of successful transfection of recombinant shRNA molecules, DNA extracted from the harvested hepatocytes was used as template for performing the PCR with specific primers for U6 Entry vector. The PCR products found at a product length of 287 bp confirmed the successful transfection of the recombinant shRNA constructs into the hepatocytes (Fig S8 & Fig S9). The total RNA isolated from the hepatocytes into which recombinant shRNA constructs were transfected, was used for the synthesis of first strand of cDNA. The cDNA was used for performing Real time PCR for assessing the expression of *SCD*, *SREBP1*and *GAPDH* genes in the hepatocytes after transfecting with recombinant shRNA constructs.

### Amplification and melting curve analysis

Amplification curves for both the target genes showed typical initiation phase, exponential and plateau phase, indicating the successful exponential amplification of the product (Fig S10). Melt curves of the amplified products displayed specific single peak for *SCD, SREBP1* and *GAPDH* genes, indicating specific amplification and homogeneity of the PCR products. These products were run on 1.5% agarose gel and expected amplicons sizes were observed.

### Silencing efficiency of anti-*SCD* and -*SREBP1* shRNA constructs

A significant (P<0.05) reduction was observed at the mRNA expression levels of *SCD* and *SREBP1* genes, after transfecting the recombinant shRNA constructs into the hepatocytes, when compared with mock control. Expression levels (40-ΔCt) of *SCD* gene after transfecting with different shRNA constructs was 37.97, 37.61, 38.76, 38.42, 39.235 and 39.41 in the cells with shRNA1, shRNA2, shRNA3, shRNA4, shRNA5 and mock control, respectively. Expression of *SCD* gene in the transfected hepatocytes with shRNA1 and shRNA2 molecules reduced significantly (P<0.05) when compared with the mock control. Knock down efficiency of shRNA1, shRNA2, shRNA3, shRNA4 and shRNA5 molecules were 66.90%, 74.21%, 42.76%, 54.78% and 20.45%, respectively. The shRNA2 molecule was found as the best molecule among all the shRNA molecules used for silencing the *SCD* gene expression *in vitro*.

In case of *SREBP1* gene, mRNA expression levels (40-ΔCt) were 29.58, 33.60, 32.31, 33.72 and 33.64 after transfecting with shRNA1, shRNA2, shRNA3, shRNA4 and shRNA5 molecules into embryonic hepatocytes, respectively. Expression levels of *SREBP1* gene in hepatocytes into which shRNA1, shRNA2, shRNA3 shRNA4 and shRNA5 molecules were transfected, reduced significantly (P<0.05) when compared with mock control. Knock down efficiency of shRNA1, shRNA2, shRNA3, shRNA4 and shRNA5 molecules were 95.85%, %, 72.36%, 26.80% and 30.50% respectively. The shRNA1 molecule was found as the best among all the shRNA molecules used for silencing the *SREBP1* gene under cell culture system. However, Western blot analysis also revealed the expression of *SCD* and *SREBP1* proteins in both transfected as well as non-transfected mock control (Fig 5 and Fig 6).

**Fig 5.**
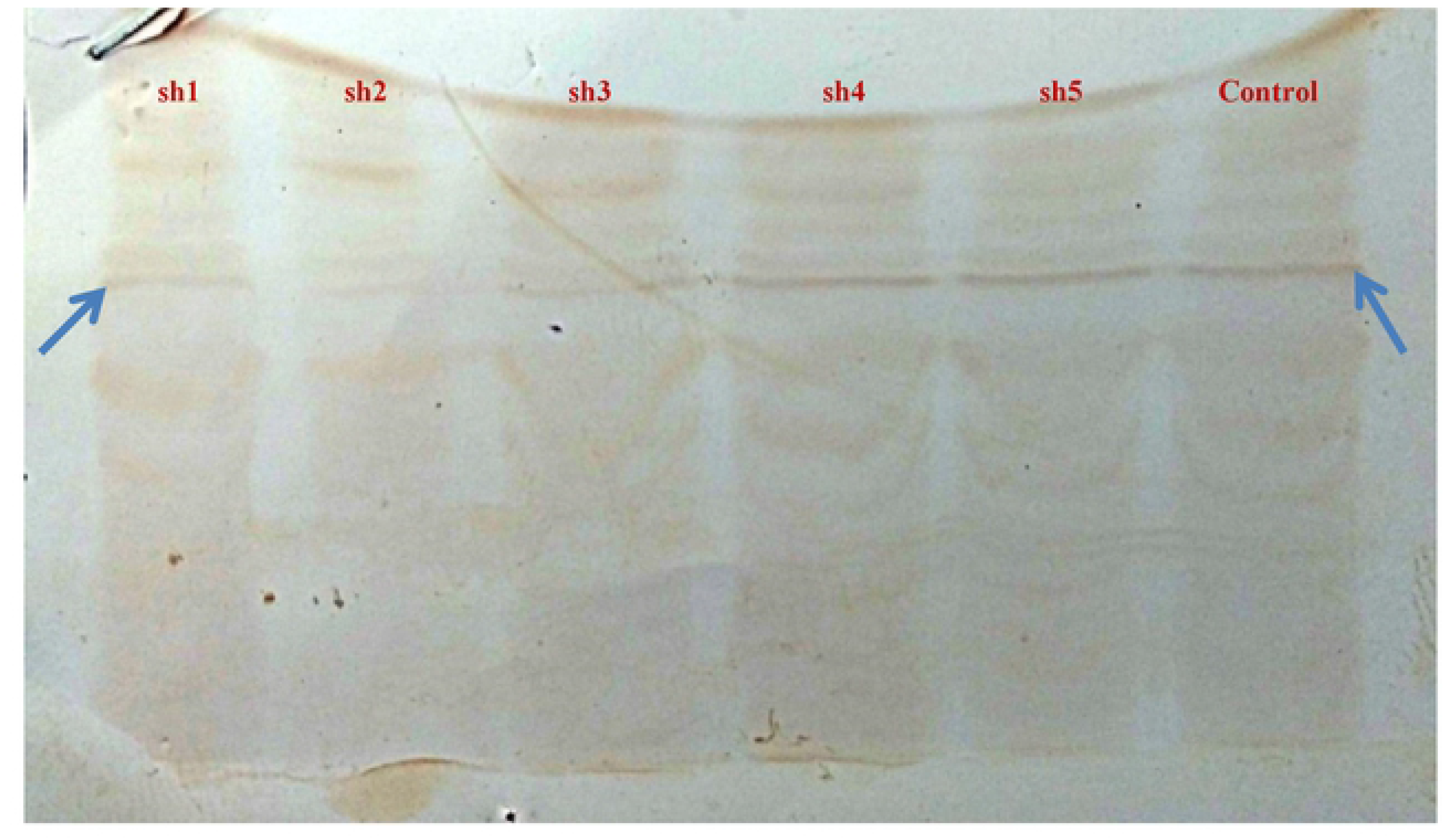
Western blot analysis of *SCD* gene silencing in transiently transfected chicken embryo hepatocytes.

**Fig 6.**
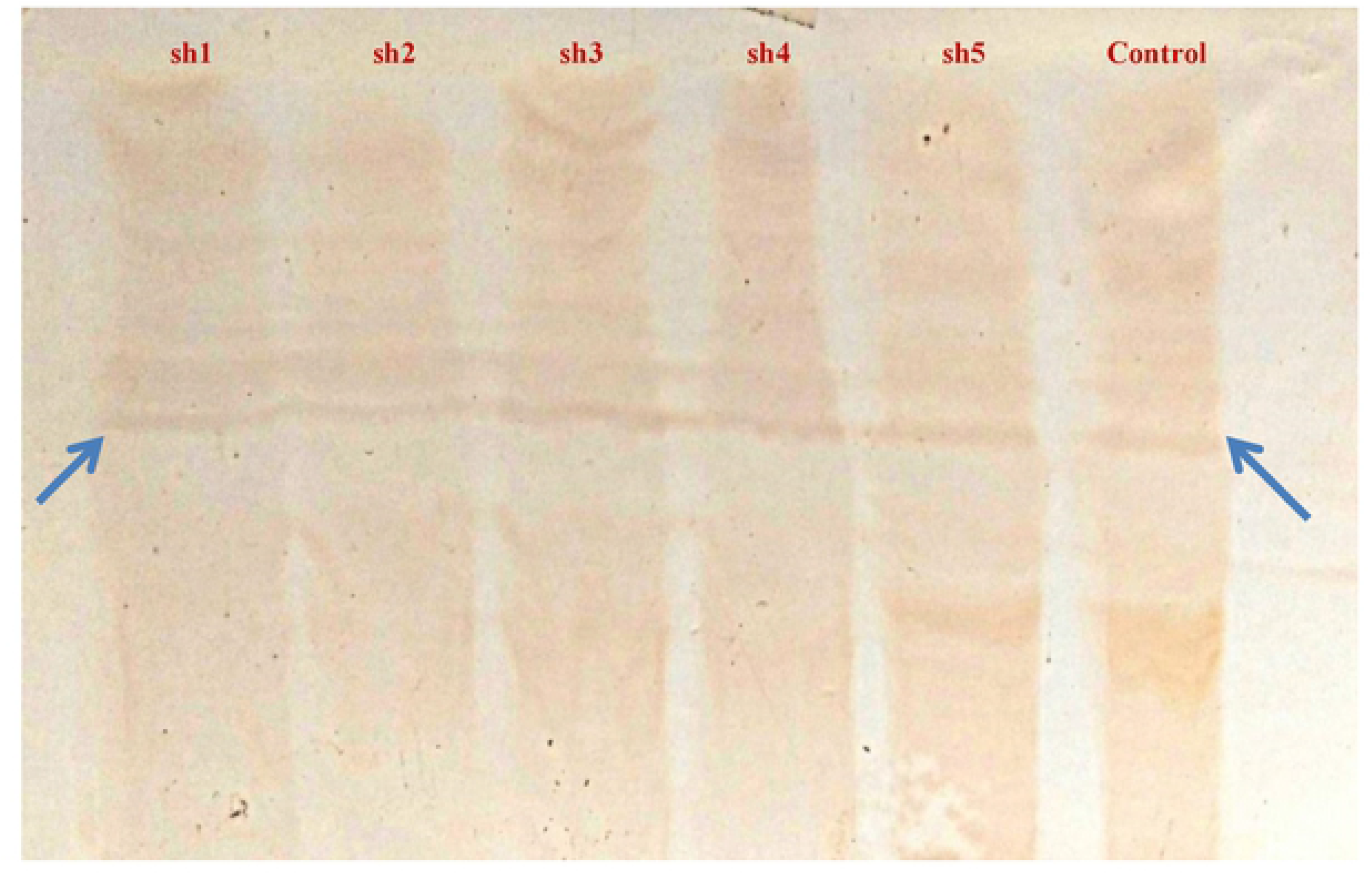
Western blot analysis of *SREBP1* gene silencing in transiently transfected chicken embryo hepatocytes.

### Relative quantification of immune response genes due to shRNA

The relative expression of immune response genes (*IFNA* and *INFB*) was monitored by quantitative PCR, after transfecting the anti-*SCD* and anti-*SREBP1* shRNA constructs into the hepatocytes. Gene expression levels of *IFNA* (40-ΔCt) was 38.525, 37.93, 37.355, 36.57, 41.23 and 34.015, while that of *IFNB* was 36.385, 35.61, 34.755, 34.815, 39.64 and 33.04 in anti-*SCD* shRNA constructs viz. shRNA1, shRNA2, shRNA3, shRNA4, shRNA5 and Scrambled shRNA, respectively (Fig 7). In case of anti-*SREBP1* shRNA constructs, the mRNA levels of *IFNA* was 35.44, 39.075, 40.2, 39.615, 40.115 and 38.23 while that of *IFNB* was 34.28, 37.5, 38.55, 38.185, 38.42 and 37.35 in the hepatocyte cells transfected with shRNA1, shRNA2, shRNA3, shRNA4, shRNA5 and Scrambled shRNA, respectively (Fig 8). There was no significant (P<0.05) difference noticed in the expression of immune response genes when compared between shRNA transfected cells with the scrambled shRNA.

**Fig 7.**
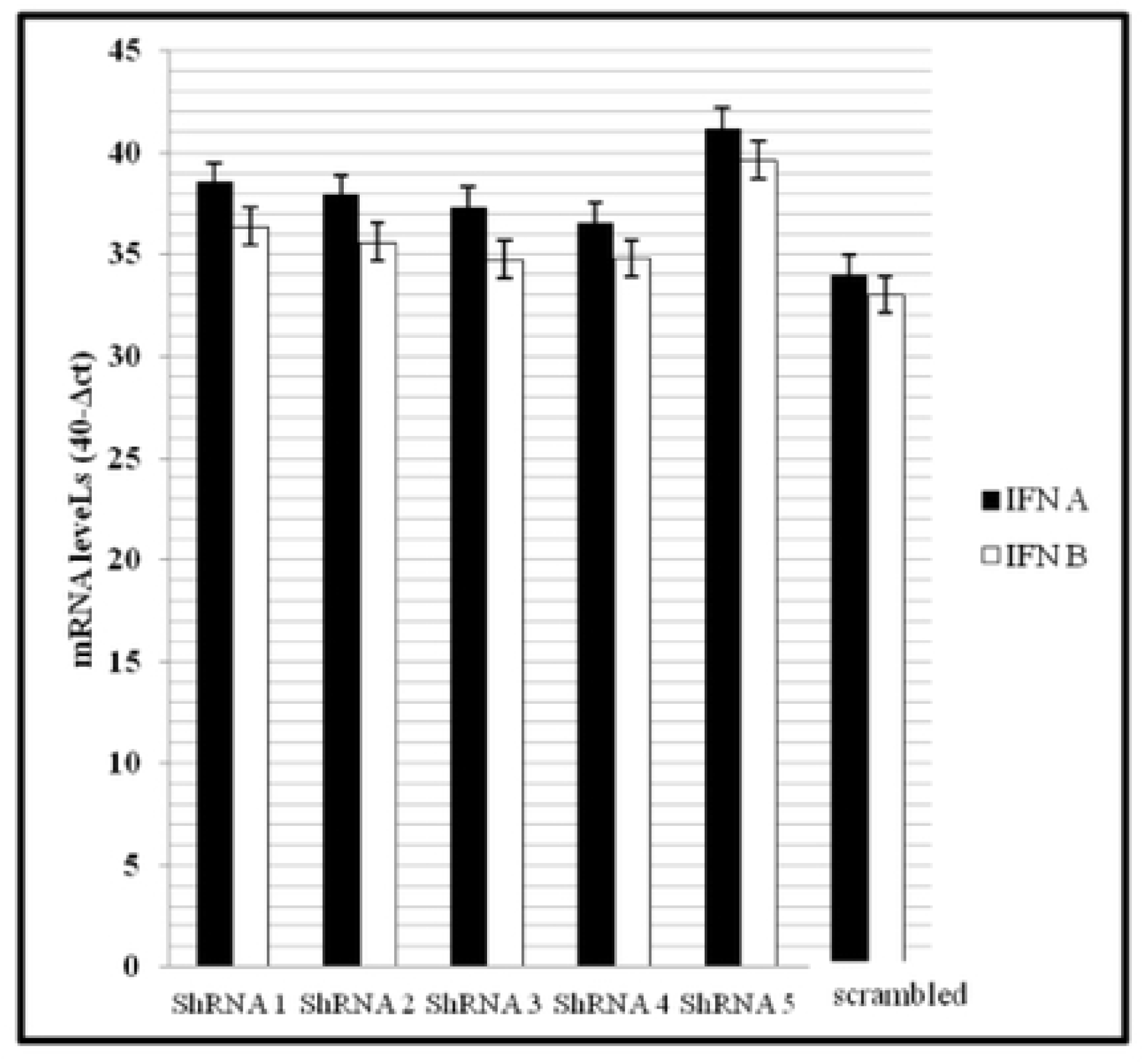
Expression of immune responsive genes in hepatocytes transfected with anti-*SCD* shRNA constructs.

**Fig 8.**
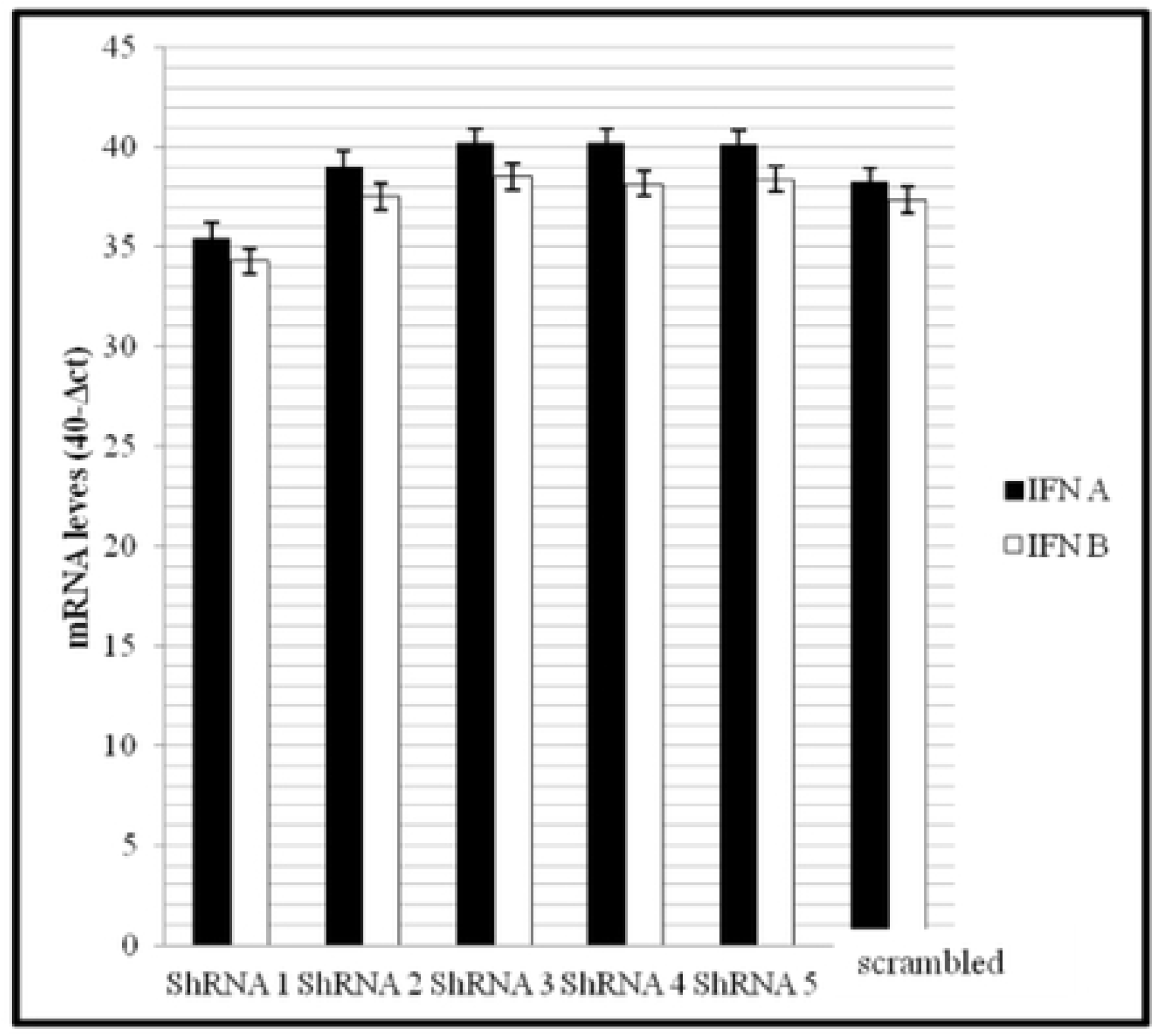
Expression of immune responsive genes in hepatocytes transfected with anti-*SREBP1* shRNA constructs.

### Sandwich ELISA for detection of *SCD* and *SREBP1* protein in cell culture

The sandwich ELISA results revealed that, the colour reaction was observed at a titre of 1:250,1:100,1:500,1:500,1:750 and 1:1000 in transfected cells with anti-*SCD* shRNA1, shRNA2, shRNA3, shRNA4, shRNA5 and scrambled shRNA. In case of anti-*SREBP1* shRNA1, shRNA2, shRNA3, shRNA4, shRNA5 and scrambled shRNA, the titre was 1:10, 1:500, 1:100, 1:500, 1:500 and 1:1000, respectively. As shRNA molecules efficiently reduced the production of *SCD* and *SREBP1* protein synthesis, those molecules which have the higher knock down efficiency (shRNA2 for *SCD*; shRNA1 and for *SREBP1*) exhibited colour reaction at lower dilutions of protein.

## Discussion

### Designed shRNA constructs against *SCD* and *SREBP1* mRNAs

RNA interference (RNAi) has become one of the powerful tools in recent times to suppress the expression of specific gene of interest [11,12], as a therapeutic option in disease management [13–16], as cancer treatment [17–19] and for understanding regulatory function of a gene by dsRNA molecules [20]. Suppression of gene expression achieved by cleavage of dsRNA molecules by ribonuclease protein (Dicer) followed by loading of siRNA molecules into RISC (RNA induced silencing complex) and then, guide strand of the siRNA molecule would guide the RISC towards the target mRNA sequence complementary with the siRNA for cleavage of target mRNA [21–25]. For successful reduction of gene expression through RNAi, designing of shRNA molecules against target gene plays crucial role [26]. In our study, anti shRNA molecules against *SCD* and *SREBP1* genes were synthesized following the criteria such as moderate to low GC content, low internal stability of sense strand at 3’ end (at least 3 bases of A/U at 15-19 positions of sense strand) and lack of internal repeats [27]. Basic Local Alignment Search Tool (BLAST) was used to detect whether our designed shRNA molecules are having the homology with any other genes to avoid the potential off-target effects.

Earlier studies revealed that the formation of secondary structures in the anti-sense/guide strand of shRNA molecule had critical role in determining the silencing efficiency and stated that a strong inverse correlation was observed between the degree of formation guide-RNA secondary structure formation and gene silencing efficiency of shRNA [28]. Patzel *et al*. (2005) classified and reported the secondary structures of guide strands of shRNA as: best silencing exhibited by sequences without any secondary structures, second best were stem loop structures with ≥ 2 free nucleotides at 5’ and ≥ 4 free nucleotides at 3’ end, next best were sequences having internal loop and two stem loop structures followed by stem loop structures with short free ends [28]. In accordance with the previous findings in our study, the shRNA molecules with best silencing efficiency (i,e shRNA2 molecule for *SCD* gene and shRNA1 molecule for *SREBP1* gene) did not posses any secondary structure in their anti-sense/guide strands. Second best shRNA molecules (i.e shRNA1 molecule for *SCD* gene and shRNA3 molecule for *SREBP1* gene) have loop structures with ≥ 2 free nucleotides at 5’ and ≥ 4 free nucleotides at 3’ end. Some of the previous studies in chicken also reported that shRNA molecules having no secondary structures in their anti-sense/guide strand had expressed high knock down efficiency in other genes such as myostatin gene and ActRIIB gene [26, 29, 30].

### Secondary structures in the target mRNA region of *SCD* and *SREBP1* gene

Earlier studies suggested that the local structure of mRNA is one of the key factors and had a strong effect on the silencing efficiency of the shRNA molecule [31–36]. All the mRNA target regions of anti-*SCD* and anti-*SREBP1* shRNA molecules revealed stem loop structures. Some studies inferred that presence of loop structures in the target mRNA regions provide easy access to the guide strand to bind with target region and had positive correlation with gene silencing efficiency [35, 37]. Presence of paired nucleotides and hairpins in the secondary structure of target mRNA region likely to have negative effect on shRNA silencing [31, 35]. In our study even though all target mRNA regions of *SCD* and *SREBP1* genes have accessibility, the differences in the silencing efficiency might be because of other responsible factor.

GC content of the target mRNA region have the crucial role in determining the silencing efficiency and it influences the loading of shRNA molecule into RISC and target affinity and specificity [10]. A high GC content in shRNA molecule hinders the dissociation of siRNA duplex inhibiting RISC loading [38, 39]. In contrary to that, some reports stated that a low GC content may reduce silencing efficiency by low target affinity and specificity [40]. The GC content of target mRNA region of our target genes ranged from 43% to 52%. Previous studies also suggested that GC content between 30% and 55% had positive correlation with silencing [27, 41, 42].

### Thermodynamic properties of shRNAs

The shRNA molecules possessing high knock down efficiency must have higher negative value of ΔG overall, ΔG duplex and lesser negative value of ΔG break-target (disruption energy) [10, 26]. Some studies postulated that shRNA molecules having the values of ΔG overall, ΔG duplex and ΔG break-target (disruption energy) within the range of - 25 to – 35 kcal/mol, - 30 to - 40 kcal/mol and - 0.5 to - 1.5 kcal/mol exhibited good silencing efficiency [26, 43]. In our study, all the shRNA molecules designed against *SCD* and *SREBP1* mRNA possessing the values of ΔG overall, ΔG duplex and ΔG break-target (disruption energy) within the desirable range have shown good silencing efficiency. The shRNA2 molecule found as the best in silencing the expression of *SCD* mRNA having higher negative value of ΔG overall, ΔG duplex and lesser negative value of ΔG break-target (disruption energy) among all shRNAs designed. Similar findings were observed in case of shRNA1 molecule, which was found as the best one in silencing the expression of *SREBP1* gene expression. Possessing the thermodynamic properties within the desirable range might be the one of the reason for better silencing efficiency of shRNA2 in *SCD* and shRNA1 in *SREBP1* genes.

### Knockdown of *SCD* and *SREBP1* expression

Silencing of the genes which are involved in the fatty acid biosynthesis has benefits like production of lean meat and egg and reduction of fat deposition in several organs of the birds. Silencing of target genes which are involved in the fat biosynthesis by using RNA interference in chicken primary hepatocyte culture helped in the devising suitable *in vitro* models for knock down of the target gene. This technique can be used further for production of the knock down chicken with reduced fat production. Till date, there were no reports available in the literature regarding the silencing of fatty acid synthesis genes in chicken. In the present study, the shRNA molecules were successfully transfected into the chicken hepatic cells and significant down regulation of the target genes were observed. These results suggest that the chicken embryo hepatocytes can be used for *in vitro* model for various functional studies. Significant reduction in the expression of *SCD* and *SREBP1* genes in hepatocytes was observed after transfecting the shRNA molecules into them. The shRNA constructs against *SCD* gene showed the knock down efficiency ranged from 20.4% to 74.2%. In case of shRNA constructs against *SREBP1* gene, they showed knock down efficiency ranging from 26.8% to 95.85%. The reasons behind the high knock down efficiency of shRNA2 against *SCD* gene and shRNA1 molecule against *SREBP1* gene might be lack of secondary structures in their anti-sense/guide strand, desirable GC percentage (43%-52%) in sense strand, more accessibility in the target mRNA region i.e stem loop structures in secondary structure of mRNA region and possessing thermodynamic properties falling in desirable range. The expression of the SCD and SREBP1 examined in the cell lysates showed lower expression in the cells transfected with shRNAs indicating the similar trends as of mRNA expression in the cell culture. Thus, these results clearly demonstrated the successful down regulation of the gene expression by designed shRNA molecules against both the target genes under *in vitro* condition. In previous reports, by using shRNA molecules in Myostatin gene [44], *ACTRIIA* gene [45] and *ACTRIIB* gene [26] knock down efficiency of 68%, 87% and 82% were found respectively in chicken embryo fibroblasts cells. In duck embryo fibroblasts, different shRNA molecules mediated through lenti-virus have down regulated the *MSTN* gene expression by 61.6, 76.9 and 79.1%, respectively, when compared with control cells [46]. In caprine foetal fibroblasts, mRNA expression levels of myostatin gene were down regulated by 89% [47] and 72% [48] after transient transfection of shRNA molecules into them.

RNA interference is powerful technique for down regulating the particular gene but it may cause the activation of immune response genes under *in vitro* and *in vivo* condition. According to the recent reports, both immune cells and non-immune cells can detect the shRNA molecules irrespective of their sequence and leads to the stimulation of interferon and inflammatory cytokines under *in vivo* and *in vitro* conditions [49, 50]. It was also reported that, shRNA molecules shorter than size of 30 bp can escape the PKR activation. It is known that the IFN response caused by activation of protein kinase R (PKR), ultimately leads to the inhibition of protein synthesis [50]. In the present study, we observed the expression of immune response genes particularly *IFNA* and *IFNB* in both tranfected and non-transfected control cells in case of both *SCD* and *SREBP1* experiments. In both the genes, the non-significant differences of expression of *IFNA* and *IFNB* genes between shRNA transfected cells and either scrambled shRNA transfected cells or un-transfected negative control embryonic hepatocyte cells. In supporting to the results obtained in the present study, Patel *et al.* (2014) and Guru Vishnu et al. (2019) in *ACTIIB* gene in goat and chicken fibroblast cells, respectively [26,51]. We suggest that shRNAs designed in our study against both *SCD* and *SREBP1* genes had the potential to be excellent shRNA molecules for further development of knock down chicken as the shRNA molecules having very good knock down efficiency escaped the interferon system of the cells.

## Conclusion

From the present findings, it is concluded that, secondary structures in the anti-sense/guide strand, desirable GC percentage in sense strand, accessibility of target mRNA region and thermodynamic properties have crucial role in deciding the knock down efficiency of shRNA molecules. Further, Chicken primary liver cell culture has been optimized and shRNA based silencing of *SCD* and *SREBP1* genes under *in vitro* cell culture system have been developed. The potential shRNA molecules (shRNA2 molecule for *SCD* and shRNA1 molecule for *SREBP1*) have been identified and can be used further *in vivo* for silencing the expression of *SCD* and *SREBP1* gene for production of the knock down chicken with possible low levels of body lipids.

## Acknowledgements

The authors convey thanks to the Indian Council of Agricultural Research, Govt. of India for financial support to carry out this research work under the National Fellow project awarded to the Corresponding author. The work was the part of the Ph.D thesis work of the first author.

## References

1. Enoch HG, Catala A, Strittmatter P. Mechanism of rat liver microsomal stearyl-CoA desaturase. Studies of the substrate specificity, enzyme-substrate interactions, and the function of lipid. Journal of Biological Chemistry. 1976 Aug 25;251(16):5095–103. PMID: 8453

2. Peláez R, Pariente A, Pérez-Sala Á, Larráyoz IM. Sterculic Acid: The Mechanisms of Action beyond Stearoyl-CoA Desaturase Inhibition and Therapeutic Opportunities in Human Diseases. Cells. 2020 Jan; 9(1):140. https://doi.org/10.3390/cells9010140 PMID: 31936134

3. Seppälä-Lindroos A, Vehkavaara S, Häkkinen AM, Goto T, Westerbacka J, Sovijärvi A, Halavaara J, Yki-Järvinen H. Fat accumulation in the liver is associated with defects in insulin suppression of glucose production and serum free fatty acids independent of obesity in normal men. The Journal of Clinical Endocrinology & Metabolism. 2002 Jul 1;87(7):3023–8. https://doi.org/10.1210/jcem.87.7.8638 PMID: 12107194

4. Brown WR, Hubbard SJ, Tickle C, Wilson SA. The chicken as a model for large-scale analysis of vertebrate gene function. Nature Reviews Genetics. 2003 Feb;4(2):87–98. https://doi.org/10.1038/nrg998 PMID: 12560806

5. Bertolio R, Napoletano F, Mano M, Maurer-Stroh S, Fantuz M, Zannini A, Bicciato S, Sorrentino G, Del Sal G. Sterol regulatory element binding protein 1 couples mechanical cues and lipid metabolism. Nature communications. 2019 Mar 22;10(1):1–1. https://doi.org/10.1038/s41467-019-09152-7 PMID: 30902980

6. Zuker M. Mfold web server for nucleic acid folding and hybridization prediction. Nucleic acids research. 2003 Jul 1;31(13):3406–15. https://doi.org/10.1093/nar/gkg595 PMID: 12824337

7. Lorenz R, Bernhart SH, Zu Siederdissen CH, Tafer H, Flamm C, Stadler PF, Hofacker IL. ViennaRNA Package 2.0. Algorithms for molecular biology. 2011 Dec 1;6(1):26.https://doi.org/10.1186/1748-7188-6-26 PMID: 22115189

8. Mathews DH, Burkard ME, Freier SM, Wyatt JR, Turner DH. Predicting oligonucleotide affinity to nucleic acid targets. Rna. 1999 Nov; 5(11):1458–69. https://doi.org/10.1017/s1355838299991148 PMID: 10580474

9. Reuter JS, Mathews DH. RNAstructure: software for RNA secondary structure prediction and analysis. BMC bioinformatics. 2010 Dec;11(1):129. https://doi.org/10.1186/1471-2105-11-129 PMID: 20230624

10. Pascut D, Bedogni G, Tiribelli C. Silencing efficacy prediction: a retrospective study on target mRNA features. Bioscience reports. 2015 Apr 1;35(2). https://doi.org/10.1042/bsr20140147 PMID: 25702798

11. Hannon GJ. RNA interference. nature. 2002 Jul;418(6894):244–51. https://doi.org/10.1038/418244a PMID: 12110901

12. Fire A, Xu S, Montgomery MK, Kostas SA, Driver SE, Mello CC. Potent and specific genetic interference by double-stranded RNA in Caenorhabditis elegans. nature. 1998 Feb;391(6669):806–11. https://doi.org/10.1038/35888 PMID: 9486653

13. Acharya R. Prospective vaccination of COVID-19 using shRNA-plasmid-LDH nanoconjugate. Medical Hypotheses. 2020 Oct 1;143:110084. https://doi.org/10.1016/j.mehy.2020.110084 PMID: 32663741

14. Kumar S, Dey A, Yau YY, Easterling M, Sahoo L. RNA Interference: For Improving Traits and Disease Management in Plants. In Climate Change, Photosynthesis and Advanced Biofuels 2020 (pp. 339–368). Springer, Singapore. https://doi.org/10.1007/978-981-15-5228-1_14 PMID: 25332689

15. Acharya R. The recent progresses in shRNA-nanoparticle conjugate as a therapeutic approach. Materials Science and Engineering: C. 2019 Nov 1;104:109928. https://doi.org/10.1016/j.msec.2019.109928 PMID: 31500065

16. Thijssen MF, Brüggenwirth IM, Gillooly A, Khvorova A, Kowalik TF, Martins PN. Gene silencing with siRNA (RNA interference): a new therapeutic option during ex vivo machine liver perfusion preservation. Liver Transplantation. 2019 Jan;25(1):140–51. https://doi.org/10.1002/lt.25383 PMID: 30561891

17. Qiao H, Wang YF, Yuan WZ, Zhu BD, Jiang L, Guan QL. Silencing of ENO1 by shRNA inhibits the proliferation of gastric cancer cells. Technology in cancer research & treatment. 2018 Jul 9;17. https://doi.org/10.1177%2F1533033818784411 PMID: 29986635

18. Ni Q, Zhang F, Zhang Y, Zhu G, Wang Z, Teng Z, Wang C, Yung BC, Niu G, Lu G, Zhang L. In situ shRNA synthesis on DNA–polylactide nanoparticles to treat multidrug resistant breast cancer. Advanced Materials. 2018 Mar;30(10):1705737. https://doi.org/10.1002/adma.201705737 PMID: 29333658

19. Shailender G, Patanla K, Malla RR. ShRNA-mediated matrix metalloproteinase-2 gene silencing protects normal cells and sensitizes cancer cells against ionizing-radiation induced damage. Journal of Cellular Biochemistry. 2020 Feb;121(2):1332–52. https://doi.org/10.1002/jcb.29369 PMID: 31489968

20. Patel M, Peter ME. Identification of DISE-inducing shRNAs by monitoring cellular responses. Cell Cycle. 2018 Feb 16;17(4):506–14. https://doi.org/10.1080/15384101.2017.1383576 PMID: 29092660

21. Bernstein E, Caudy AA, Hammond SM, Hannon GJ. Role for a bidentate ribonuclease in the initiation step of RNA interference. Nature. 2001 Jan;409(6818):363–6. https://doi.org/10.1038/35053110 PMID: 11201747

22. Zamore PD, Tuschl T, Sharp PA, Bartel DP. RNAi: double-stranded RNA directs the ATP-dependent cleavage of mRNA at 21 to 23 nucleotide intervals. Cell. 2000 Mar 31;101(1):25–33. https://doi.org/10.1016/s0092-8674(00)80620-0 PMID: 10778853

23. Elbashir SM, Harborth J, Lendeckel W, Yalcin A, Weber K, Tuschl T. Duplexes of 21-nucleotide RNAs mediate RNA interference in cultured mammalian cells. nature. 2001 May;411(6836):494–8. https://doi.org/10.1038/35078107 PMID: 11373684

24. Hammond SM, Bernstein E, Beach D, Hannon GJ. An RNA-directed nuclease mediates post-transcriptional gene silencing in Drosophila cells. Nature. 2000 Mar;404(6775):293–6. https://doi.org/10.1038/35005107 PMID: 10749213

25. Hammond SM, Caudy AA, Hannon GJ. Post-transcriptional gene silencing by double-stranded RNA. Nature Reviews Genetics. 2001 Feb;2(2):110–9. https://doi.org/10.1038/35052556 PMID: 11253050

26. Vishnu PG, Bhattacharya TK, Bhushan B, Kumar P, Chatterjee RN, Paswan C, Dushyanth K, Divya D, Prasad AR. In silico prediction of short hairpin RNA and in vitro silencing of activin receptor type IIB in chicken embryo fibroblasts by RNA interference. Molecular biology reports. 2019 Jun 1;46(3):2947–59. https://doi.org/10.1007/s11033-019-04756-0 PMID: 30879273

27. Reynolds A, Leake D, Boese Q, Scaringe S, Marshall WS, Khvorova A. Rational siRNA design for RNA interference. Nature biotechnology. 2004 Mar;22(3):326–30. https://doi.org/10.1038/nbt936 PMID: 14758366

28. Patzel V, Rutz S, Dietrich I, Köberle C, Scheffold A, Kaufmann SH. Design of siRNAs producing unstructured guide-RNAs results in improved RNA interference efficiency. Nature biotechnology. 2005 Nov;23(11):1440–4. https://doi.org/10.1038/nbt1151 PMID: 16258545

29. Tripathi AK, Aparnathi MK, Patel AK, Joshi CG. In vitro silencing of myostatin gene by shRNAs in chicken embryonic myoblast cells. Biotechnology progress. 2013 Mar;29(2):425–31. https://doi.org/10.1002/btpr.1681 PMID: 23292805

30. Bhattacharya TK, Shukla R, Chatterjee RN, Dushyanth K. Knock down of the myostatin gene by RNA interference increased body weight in chicken. Journal of biotechnology. 2017 Jan 10;241:61–8. https://doi.org/10.1016/j.jbiotec.2016.11.012 PMID: 27845166

31. Kretschmer-Kazemi Far R, Sczakiel G. The activity of siRNA in mammalian cells is related to structural target accessibility: a comparison with antisense oligonucleotides. Nucleic acids research. 2003 Aug 1;31(15):4417–24. https://doi.org/10.1093/nar/gkg649 PMID: 12888501

32. Bohula EA, Salisbury AJ, Sohail M, Playford MP, Riedemann J, Southern EM, Macaulay VM. The efficacy of small interfering RNAs targeted to the type 1 insulin-like growth factor receptor (IGF1R) is influenced by secondary structure in the IGF1R transcript. Journal of Biological chemistry. 2003 May 2;278(18):15991–7. https://doi.org/10.1074/jbc.m300714200 PMID: 12604614

33. Stewart CK, Li J, Golovan SP. Adverse effects induced by short hairpin RNA expression in porcine fetal fibroblasts. Biochemical and biophysical research communications. 2008 May 23;370(1):113–7. https://doi.org/10.1016/j.bbrc.2008.03.041 PMID: 18358831

34. Luo KQ, Chang DC. The gene-silencing efficiency of siRNA is strongly dependent on the local structure of mRNA at the targeted region. Biochemical and biophysical research communications. 2004 May 21;318(1):303–10. https://doi.org/10.1016/j.bbrc.2004.04.027 PMID: 15110788

35. Schubert S, Grünweller A, Erdmann VA, Kurreck J. Local RNA target structure influences siRNA efficacy: systematic analysis of intentionally designed binding regions. Journal of molecular biology. 2005 May 13;348(4):883–93.https://doi.org/10.1016/j.jmb.2005.03.011 PMID: 15843020

36. Gredell JA, Berger AK, Walton SP. Impact of target mRNA structure on siRNA silencing efficiency: a large-scale study. Biotechnology and bioengineering. 2008 Jul 1;100(4):744–55. https://doi.org/10.1002/bit.21798 PMID: 18306428

37. Overhoff M, Alken M, Far RK, Lemaitre M, Lebleu B, Sczakiel G, Robbins I. Local RNA target structure influences siRNA efficacy: a systematic global analysis. Journal of molecular biology. 2005 May 13;348(4):871–81. https://doi.org/10.1016/j.jmb.2005.03.012 PMID: 15843019

38. Holen T. Mechanisms of RNAi: mRNA cleavage fragments may indicate stalled RISC. Journal of RNAi and gene silencing: an international journal of RNA and gene targeting research. 2005 Aug;1(1):21..PMID: 19771200

39. Matveeva O, Nechipurenko Y, Rossi L, Moore B, Sætrom P, Ogurtsov AY, Atkins JF, Shabalina SA. Comparison of approaches for rational siRNA design leading to a new efficient and transparent method. Nucleic acids research. 2007 Apr 1;35(8):e63. https://dx.doi.org/10.1093%2Fnar%2Fgkm088 PMID: 17426130

40. Li W, Cha L. Predicting siRNA efficiency. Cellular and molecular life sciences: CMLS. 2007 Jul;64(14):1785–92. https://doi.org/10.1007/s00018-007-7057-3 PMID: 17415516

41. Shah JK, Garner HR, White MA, Shames DS, Minna JD. sIR: siRNA Information Resource, a web-based tool for siRNA sequence design and analysis and an open access siRNA database. BMC bioinformatics. 2007 Dec 1;8(1):178. https://doi.org/10.1186/1471-2105-8-178 PMID: 17540034

42. Wang L, Mu FY. A Web-based design center for vector-based siRNA and siRNA cassette. Bioinformatics. 2004 Jul 22;20(11):1818–20. https://doi.org/10.1093/bioinformatics/bth164 PMID: 15001477

43. Shao Y, Chan CY, Maliyekkel A, Lawrence CE, Roninson IB, Ding Y. Effect of target secondary structure on RNAi efficiency. Rna. 2007 Oct 1;13(10):1631–40. https://dx.doi.org/10.1261%2Frna.546207 PMID: 17684233

44. Tripathi AK, Aparnathi MK, Vyavahare SS, Ramani UV, Rank DN, Joshi CG. Myostatin gene silencing by RNA interference in chicken embryo fibroblast cells. Journal of biotechnology. 2012 Aug 31;160(3-4):140–5. https://doi.org/10.1016/j.jbiotec.2012.03.001 PMID: 22445467

45. Satheesh P, Bhattacharya TK, Kumar P, Chatterjee RN, Dhara SK, Paswan C, Shukla R, Dushyanth K. Gene expression and silencing of activin receptor type 2A (ACVR2A) in myoblast cells of chicken. British poultry science. 2016 Nov 1;57(6):763–70. https://doi.org/10.1080/00071668.2016.1219693 PMID: 27635666

46. Tao Z, Zhu C, Song C, Song W, Ji G, Shan Y, Xu W, Li H. Lentivirus-mediated RNA interference of myostatin gene affects MyoD and Myf5 gene expression in duck embryonic myoblasts. British poultry science. 2015 Sep 3;56(5):551–8. https://doi.org/10.1080/00071668.2015.1085958 PMID: 26301941

47. Kumar R, Singh SP, Kumari P, Mitra A. Small interfering RNA (siRNA)-mediated knockdown of myostatin influences the expression of myogenic regulatory factors in caprine foetal myoblasts. Applied biochemistry and biotechnology. 2014 Feb 1;172(3):1714–24. https://doi.org/10.1007/s12010-013-0582-7 24254256

48. Jain SK, Jain H, Kumar D, Bedekar MK, Pandey AK, Sarkhel BC. Quantitative evaluation of myostatin gene in stably transfected caprine fibroblast cells by anti-myostatin shRNA. Applied biochemistry and biotechnology. 2015 Sep 1;177(2):486–97. https://doi.org/10.1007/s12010-015-1757-1 PMID: 26234434

49. Sledz CA, Holko M, De Veer MJ, Silverman RH, Williams BR. Activation of the interferon system by short-interfering RNAs. Nature cell biology. 2003 Sep;5(9):834–9. https://doi.org/10.1038/ncb1038 PMID: 12942087

50. Judge AD, Sood V, Shaw JR, Fang D, McClintock K, MacLachlan I. Sequence-dependent stimulation of the mammalian innate immune response by synthetic siRNA. Nature biotechnology. 2005 Apr;23(4):457–62. https://doi.org/10.1038/nbt1081 PMID: 15778705

51. Patel AK, Tripathi AK, Shah RK, Patel UA, Joshi CG. Assessment of goat activin receptor type IIB knockdown by short hairpin RNAs in vitro. Journal of Receptors and Signal Transduction. 2014 Dec 1;34(6):506–12. https://doi.org/10.3109/10799893.2014.922574 PMID: 24870261

